# Scalable Production of Human Erythrocytes from Induced Pluripotent Stem Cells

**DOI:** 10.1101/050021

**Authors:** Ying Wang, Yongxing Gao, Chaoxia He, Zhaohui Ye, Sharon Gerecht, Linzhao Cheng

## Abstract

In vitro production of erythrocytes in physiologic numbers from human induced pluripotent stem cells (hiPSCs) holds great promise for improved transfusion medicine and novel cell therapies. We report here, for the first time, a strategy for scalable and xeno-free differentiation of hematopoietic stem/progenitor cells from hiPSCs and subsequent erythrocytes specification, by using stepwise cell culture conditions and by integrating spinner flasks and rocker. This system supported robust and reproducible definitive hematopoietic differentiation of multiple hiPSC lines. We demonstrated an ultra-high yield of up to 4×10^9^ CD235a^+^ erythrocytes at >98% purity when using a 1-litre spinner flask for suspension culture. Erythrocytes generated from our system can reach a mature stage with red blood cell (RBC) characteristics of enucleation, β-globin protein expression and oxygen-binding ability. The entire process is xeno-free and clinically compliant, allowing future mass production of hiPSC-derived RBCs for transfusion medicine purposes.

## Introduction

As an irreplaceable clinical practice in modern medicine, red blood cell (RBC) transfusion saves lives of patients suffering from severe blood lose or chronic anemia. Accessibility and dependence on blood donation remain critical around the world. This is a particularly important issue for the patients who need RBCs of rare blood types, because of severe complications caused by mismatching transfusion or allo-sensitized responses among chronically transfusion-dependent patients^1^.

A potential solution is to manufacture RBCs from hematopoietic stem and progenitor cells (HSPCs) that are isolated from patients themselves or donors with desirable blood types. Recently, mature RBCs that were generated *ex vivo* from CD34^+^ HSPCs were transfused into a volunteer to demonstrate *in vivo* safety^2^, albeit at limited quantity (~4×10^10^ RBCs or ~2% of one transfusion unit). One major challenge for clinical applications has been how to expand HSPCs extensively because committed erythroid progenitors have limited proliferative potential before differentiating into mature RBCs^3^.

Human induced pluripotent stem cells (hiPSCs) is a promising cell source that can be easily derived from donor somatic cells and expanded indefinitely^4^. They would provide a potentially unlimited source of RBCs perfectly matching to patients or a favorable donor with any antigen types, especially the universal donor type O and Rhesus D-negative (O Rh-) and rare-phenotype matching RBCs^5-8^. Recently, hiPSCs derived from healthy donors or patients with blood-related disorders were successfully differentiated towards erythroid lineages and completed terminal maturation, which is marked by enucleation, blood antigen display, fetal-to-adult hemoglobin switch, functional oxygen carrying ability and many other characteristic features comparable with primary blood cells or those derived from primary HSPCs^9-13^.

The current methodology for the *in vitro* generation of RBCs from hiPSCs generally includes three distinguishable steps: (1) formation of definitive HSPCs that express both CD34 and CD45 markers (CD34^+^CD45^+^); (2) erythroblast specification and expansion; and (3) erythrocyte maturation. To complete step (1), various protocols were established to promote mesodermal commitment and hematopoietic specification of hiPSCs^14^. For example, co-culturing with mouse stromal cells^9,15, 16^ or human stromal cells^17-19^ harbored hiPSCs or HSPCs in stromal niches that recapitulated the *in vivo* environments of hematopoiesis, and sometimes was integrated together with the latter two steps. Efficient feeder-free differentiation of hiPSC in monolayer adhesion culture on supporting substrates has not been established until a recent study showed that tenascin C could replace confluent OP-9 mouse stromal cell line, although the erythrocyte production from the new culture system has not been demonstrated^20^. The most commonly used method for step (1) so far is by forming embryoid bodies (EBs) that are self-organized to mimic early human embryonic structures and development^21^. The mesodermal commitment and haematopoietic differentiation with EBs are further enhanced by adding exogenous cytokines such as bone morphogenetic protein 4 (BMP4), fibroblast growth factor 2 (FGF2), stem cell factor (SCF), vascular endothelial growth factor (VEGF), interleukin-3 (IL-3), etc.^12, 22-26^. There are a number of approaches to form EBs, such as hanging drop, self-aggregation in static/dynamic suspension and forced aggregation (FA) method. Among these approaches, others and we found that FA in 96-well plates and micro-patterned plates that forms individual EBs while preventing multiple EBs from fusion and agglomeration, efficiently generate CD34^+^CD45^+^ definitive HSPCs^10, 27-31^. In our most recent study, we demonstrated a FA differentiation method in 96-well plates with modified xeno-free medium to provide robust and reproducible HSPC generation from hiPSCs^30^, which can be used as a baseline for further improvement. For step (2) and (3), efficient stromal-free suspension culture was established to generate enucleated and hemoglobin-expressing erythrocytes using medium that contained erythropoietin (EPO) and human plasma^10-12^.

Considering a manufacturing process required for clinical purposes, one has to implement a culture procedure with vessels that are suitable for large-scale cell culture and expansion. Most EB differentiation strategies involve separation of individual EBs using multi-well plates. They can only accommodate a pilot scale production in a scale-out process by linearly increasing the number of culture vessels, which requires investment of advanced automation equipment if technically feasible. In contrast, the methods of forming EBs in suspension showed the capacity to adapt to vessels that were compliant with scale-up bioprocess and generate matured cells, such as cardiomyocytes, myeloid cells, and hepatocytes^26, 32-35^. In these studies, spinner flasks and flat tissue culture flasks placed on a rocker were commonly used due to the scalability of directly transferring to well-established stirred tank bioreactors (up to 2000 L) and wave bag bioreactors (up to 1000 L), respectively. These bioreactor systems are widely utilized in industrial biotechnology for multi-parametric monitoring/control. However, this kind of volumetrically scalable system for clinical manufacturing purpose has not yet been established for the generation of human erythrocytes from hiPSCs.

In this study, we aim to establish a scalable system that can support large-scale EB differentiation towards hematopoietic lineages and high-yield erythrocyte production from hiPSCs *in vitro*. To achieve this goal, we first sought to combine the expansion of undifferentiated hiPSCs and the formation of EBs in suspension using spinner flasks. Then we integrated this process with static and dynamic culture on a platform rocker to enhance the efficiency of HSPC generation. HSPCs differentiated in the scalable system were directly induced to generate mature erythrocytes in suspension in scalable settings.

## Results

### E8-based Medium in Spinner Flasks is a Superior Condition for Initiating Scalable EB Formation

Utilizing our scalable expansion of undifferentiated hiPSC in suspension, which is matrix-independent and microcarrier-free^36^, we transitioned to volumetrically scalable EB differentiation using vessels such as a spinner flask (SF) and a tissue culture flask on a platform rocker (TR). A culture in spinner flasks can be directly scale-up into large stirred tank bioreactors, whereas a culture in TR condition is suitable for straightforward scale-up into a WAVE bag bioreactor. Shear stress on hiPSCs in the suspension culture in agitating vessels is known to affect cell survival^37^. Medium flow in a SF and TR generate different levels of shear stress. Thus we first tested which scalable system better support EB formation from a single cell suspension from BC1 hiPSCs^38^. We found that the serum-free medium (SFM) used in the standard forced aggregation (FA) differentiation for the first 2 days^30^ could not support consistent bulk EB formation in suspension culture especially in spinner flasks, resulting in an unexpected high level of cell death with cell viability below 65% (Fig. 1a). In contrast, EBs formed with a modified E8-based medium supplemented with human albumin and the same amount of BMP4 and a ROCK inhibitor Y27632 as the SFM (called E8-EB medium thereafter) showed significantly higher cell viability, 89±4 % on EB day 1 (Fig. 1a), while with similar total cell yield (data not shown). It demonstrated a simple but efficacious approach to reduce cell death during EB formation in spinner flasks causing by shear force. We also found that EBs formed in the E8-EB medium in spinner flasks were more uniform in size, while many large EBs with irregular shapes formed on rocker likely due to insufficient and uneven agitation (Fig. 1b). Quantitative analysis of EB size indicated that EBs formed in spinner flasks have a homogeneous size with an average equivalent diameter of 187±36 μm, whereas a wider distribution of EB size was found on rocker (247±128 μm) with many compact EBs larger than 400 μm (Fig. 1c), which would cause cell death in the center due to insufficient transport of nutrient, waste and O_2_. Therefore, we chose to use E8-EB medium in spinner flasks as the optimized condition to form spatially and temporally synchronized EBs.

**Figure 1.**
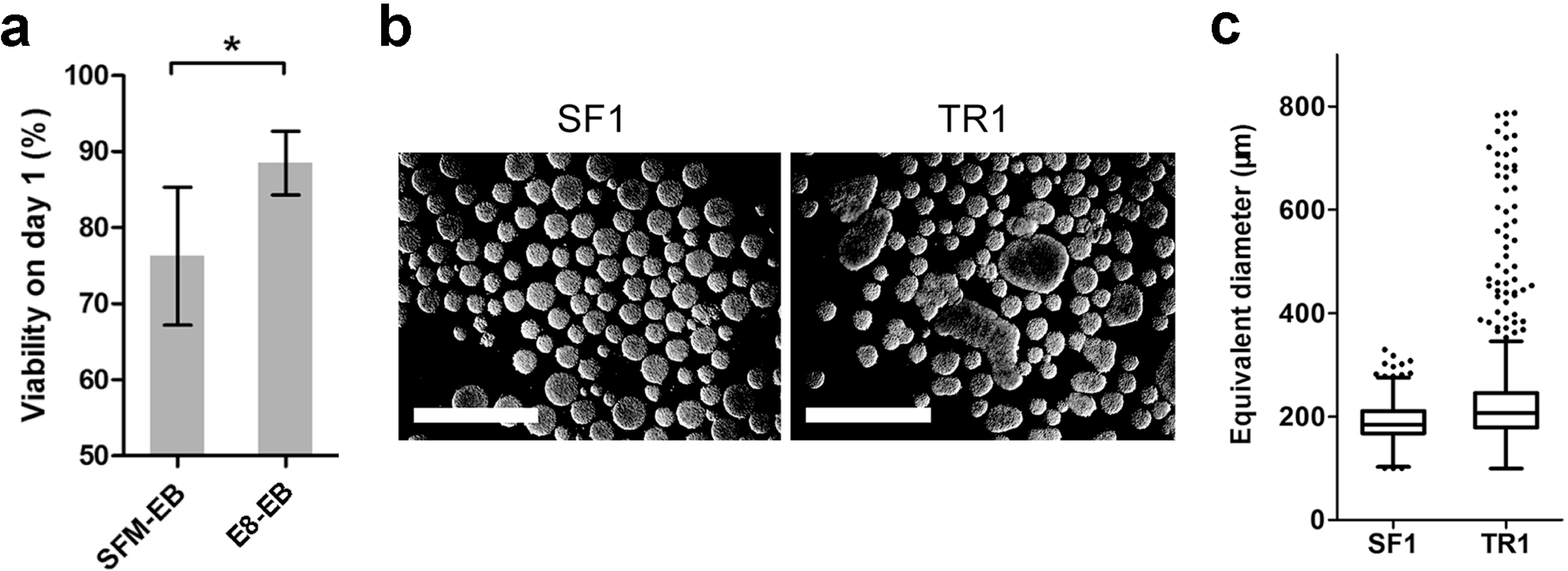
Optimization of initial EB formation in suspension cultures. (a) Levels of cell viability on day 1 using different media for EB formation in suspension cultures. All media were supplemented with 10 μM Y27632 to promote cell survival. SFM-EB: serum-free medium with BMP4 and FGF2 as the medium used in the FA control of EB formation; E8-EB: E8 medium with BMP4 and human albumin (n = 3 independent experiments, mean ± s.d.). (b) Representative images of EBs formed at day 1 in spinner flask at 35 rpm (SF1) and 150-cm^2^ tissue culture flask on rocker at 10 rpm on day 1 (TR1), respectively. Irregular shape and sizes were observed in TR day 1 EBs (scale bars = 1 mm). (c) Size of EBs at day 1 generated in spinner flask (SF1) or on rocker (TR1). More than 450 EBs in each condition from three independent experiments were analysed and illustrated in Whiskers-Tukey plot (n = 3 independent experiments, mean ± s.e.m.).

### An Optimized integration of Static Culture and Spinner Flasks Facilitates Scalable Generation of Hematopoietic Cells

The yield from one bioreactor run (indicated as “batch yield”) is an essential parameter to measure outcomes in the scale-up process. In order to maximize the yield of the EB differentiation, we explored several strategies using SF and TR (Fig. 2a), based on media and timeline modified from the FA method previously established^30^. After formed in spinner flasks, EBs harvested on day 3 were cultured either in spinner flasks or on rocker. In one condition, a 3-day static culture in tissue culture flasks (TS, see illustration in Fig. 2a) was also applied between the spinner and the rocking condition (SF-TS-TR). We compared the yield of suspension cells released from EBs (harvested at day 13) in batches with 40-ml initial volume followed by different combinations of these module conditions (Fig. 2a). From four independent experiments in each condition, the conventional FA method generated 10.2±4.2×10^6^ live cells from eight 96-well plates starting from 40-ml of initial single cell suspension (Fig. 2b). With the same initial volume, the methods in which EBs were in free suspension throughout the culture (such as SF, TR and SF-TR, see Fig. 2a) showed poor reproducibility with about 50% of experiments failing to generate hematopoietic cells. Only the combinatorial SF-TS-TR strategy including a 3-day static culture in tissue culture flasks supported more consistent differentiation with higher yields. A total of 29.1±10.8 ×10^6^ live cells were generated from SF-TS-TR suspension, 2.9-fold higher than the FA control (Fig. 2b).

**Figure 2.**
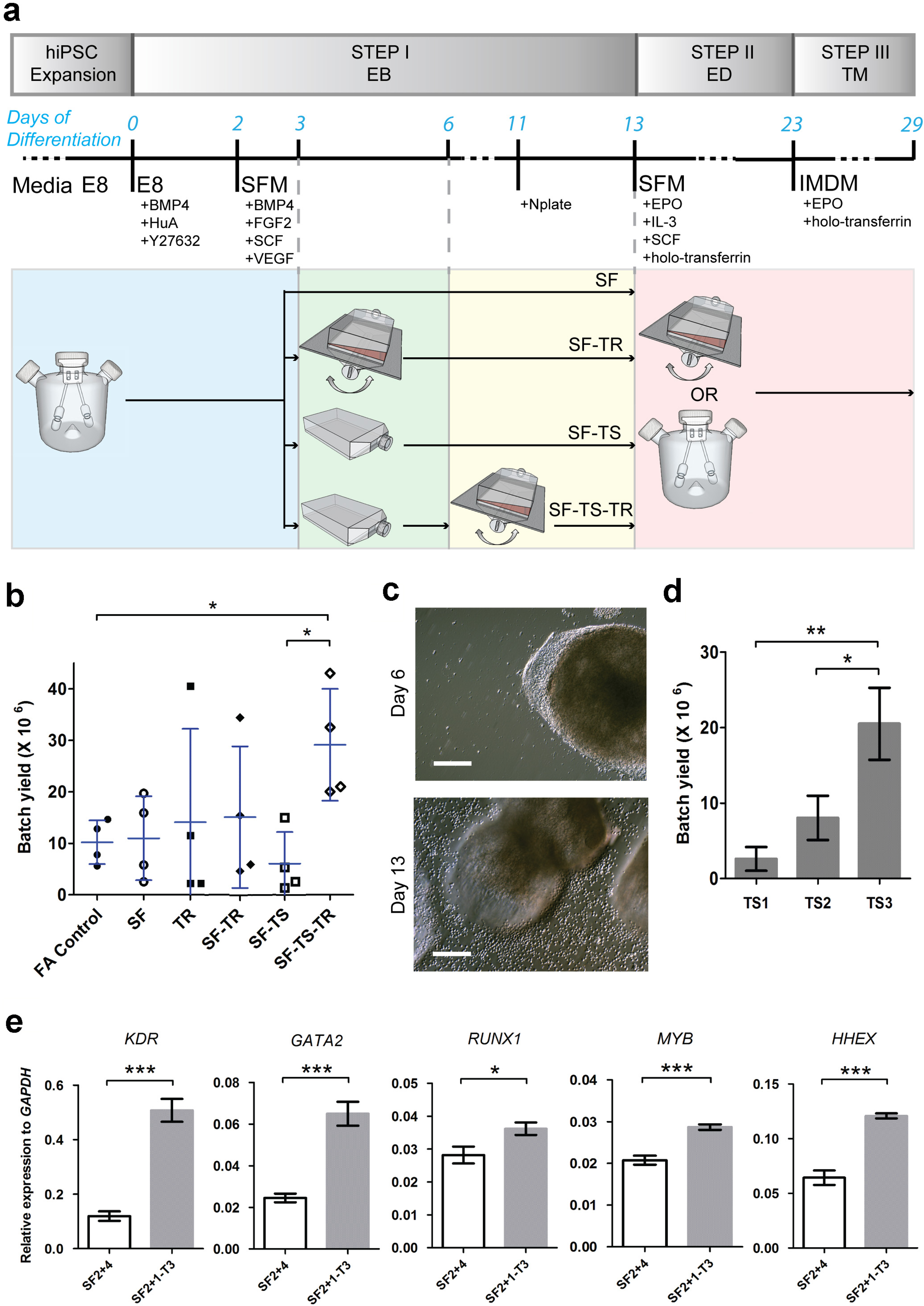
Establishment and optimization of a scalable protocol for hematopoietic cell generation in suspension. (a) Schematic presentation of strategies for scalable EB differentiation and production of hiPSC-derived erythrocytes. The differentiation was induced in three successive steps: embryoid body differentiation (EB), erythroid differentiation (ED) and terminal maturation (TM). Media and culture condition changes are illustrated according to the timeline of differentiation. (b) Yield of single suspension cells on day 13 from one batch using different EB differentiation protocols. SF and TR represent continuous EB differentiation for 13 days in spinner flasks and tissue culture flasks on rocker, respectively. SF-TS or SF-TR represent additional 8 days in static or rocking condition, respectively, after 3-day EB formation in spinner flasks (n = 4 independent experiments; mean ± s.e.m.). (c) Exemplary light microscopy images of EB differentiation using SF-TS-TR on day 6 and day 13. EB attachment on day 6 and single hematopoietic suspension cells on day 13 were clearly shown (scale bars = 200 μm). (d) Batch yield of total single suspension cells from 40-ml cultures using different SF-TS-TR strategies with static culture for 1 day (TS1), 2 days (TS2) and 3 days (TS3) (n = 3 independent experiments; mean ± s.d.). (e) Expression of *KDR*, *GATA2*, *RUNX1*, *MYB* and *HHEX* genes in EB cells from continuous agitating SF or SF-TS strategies on EB day 6 (normalized to endogenous *GAPDH* as a housekeeping control; n = 2 independent experiments, mean ± s.d.).

Next, we investigated the potential mechanisms underlying the improvement in yield when using SF-TS-TR strategy. The three days of static culture was the only difference between SF-TS-TR and SF-TR. Light microscopy images revealed distinguishable changes of EB morphology and confirmed the emergence of HSPC on day 13 (Supplementary Fig. S1 and Fig. 2c). Under the FA condition, EBs formed after 2-3 days of centrifugal aggregation appeared loosely attaching to the bottom of the well (even in low-attachment or untreated plates). Cystic structure inside EBs emerged on or around day 5, indicating the embryonic development and cell differentiation (Supplementary Fig. S1). In SF-TS-TR method, hiPSCs formed more homogenous EBs in size on day 2 compared to FA or SF-TR (Supplementary Fig. S1). Three days in static condition provided the environment for the EBs to loosely attach to the bottom of tissue culture flasks (Fig. 2c), similar to the FA method compared to completely free floating EBs in SF-TR. It was clear that the three days of static culture was beneficial for a high yield of hematopoietic cells in suspension on day 13 when compared to 2-day or 1-day static culture (Fig. 2d). After three days in static condition, genes associated with mesoderm commitment and hemato-vascular lineages were all elevated (Supplementary Fig. S2). As a result, cells through the 3-day static condition showed significantly higher expression of key genes related to mesoderm formation (*KDR*)^16^, hemato-vascular development (*GATA2 and RUNX1*)^20, 39^ and definitive hematopoiesis (*MYB* and *HHEX*)^40-42^ compared to cells from suspension EB differentiation in spinner flasks (Fig. 2e). This is in line with the higher hematopoietic yield from cultures with 3-day in static condition in comparison to 1-day or 2-day in static condition (Fig. 2d).

### Single Cells from Scalable EB Differentiation Are Characterized as HSPCs

Recently, we showed that the addition of 20 ng/ml Romiplostium (a thrombopoietin or TPO analog commercialized as Nplate®) during the last stage of EB-mediated differentiation (day 11 to day 14) increased CD34^+^CD45^+^ HSPCs yield using the conventional FA method, and also enhanced subsequent megakaryocytic differentiation as a replacement of TPO^30^. In the current study, we also found that adding Nplate on day 11 to the SF-TS-TF culture also increased the numbers of total and CD34^+^CD45^+^ HSPCs of suspension cells harvested at day 13-14 (Supplementary Fig. S3). Therefore, we included the Nplate treatment (one dose added at day 11) in the combinatorial SF-TS-TR strategy as well as all the FA control experiments (Fig. 2a).

Similar to the FA control, suspension cells harvested from SF-TS-TR at EB day 13 contained 75.6±12.3 % CD34^+^ cells and 64.2±11.9 % CD34^+^CD45^+^ cells, with little erythrocytes that should express CD235a (Fig. 3a). The batch yield of CD34^+^CD45^+^ cells from SF-TS-TR was 14.6±4.5×10^6^, which was significantly higher than the non-scalable FA method (Fig. 3b). Colony-forming unit (CFU) assay was performed to test the level of hematopoietic progenitor cells in the total and CD34^+^ purified suspension cell populations (Fig. 3c). These results indicated that the CD34^+^CD45^+^ cells generated from SF-TS-TR had comparable contents of myeloid and erythroid progenitor cells as the FA method.

**Figure 3.**
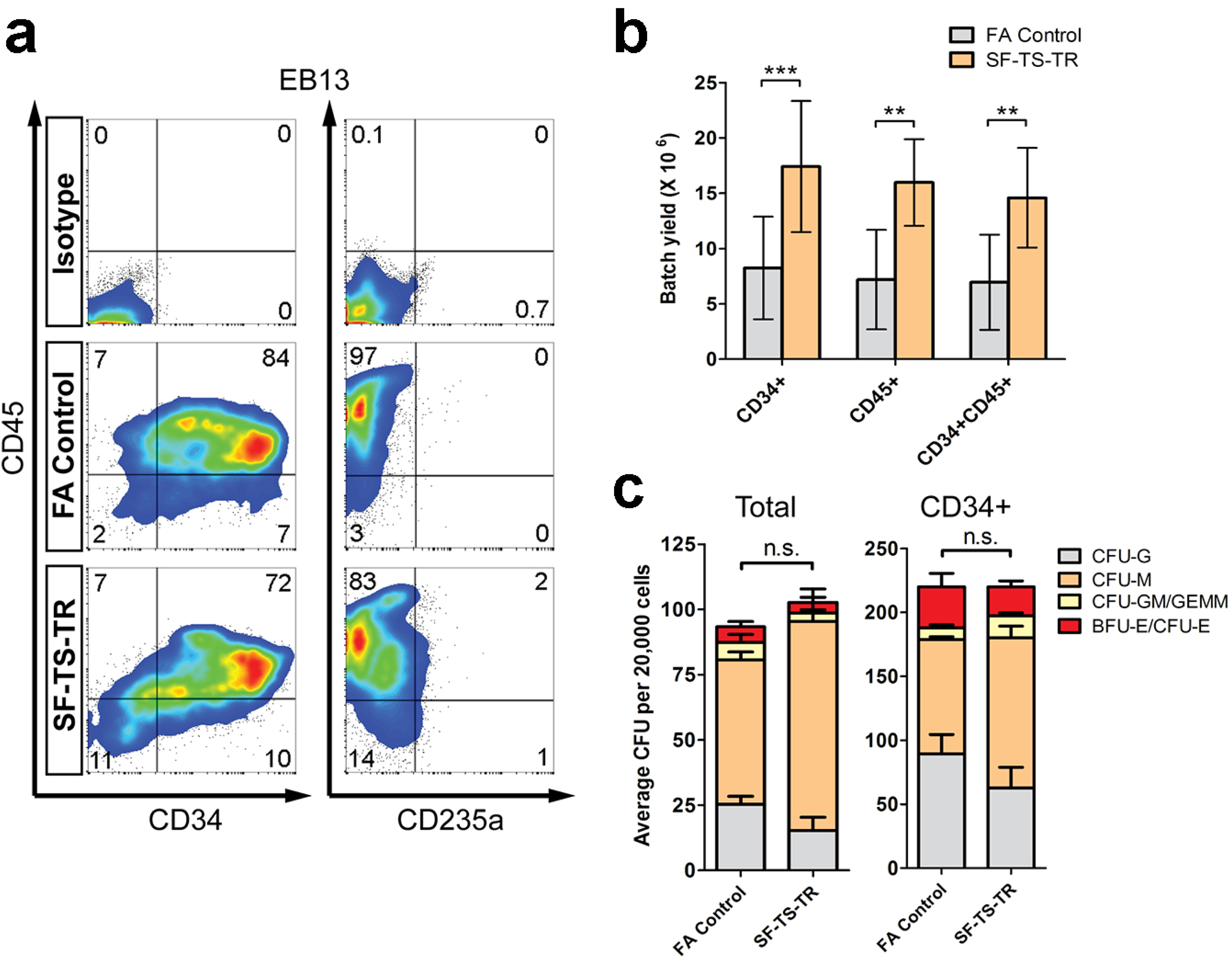
Characterizations of differentiated hematopoietic cells derived from hiPSCs using the SF-TS-TR protocol. (a) Representative flow cytometric analysis of CD34, CD45, and CD235a in the suspension cell populations harvested on EB day 13. Gates were set based on corresponding isotype controls. (b) Quantification of the batch yield of CD34^+^, CD45^+^, and CD34^+^CD45^+^ cells from SF-TS-TR method (n = 4 independent experiments, mean ± s.d.). (c) CFU assays of total suspension cells and anti-human CD34-bead isolated suspension single cells from SF-TS-TR method. After isolation, >95% of the cells showed CD34^+^ by flow cytometric analysis. The total number of CFU was analysed by the two-tailed Student’s t-test (n = 2 independent experiments, mean ± s.d.). CFU-G: colony-forming unit-granulocyte; M: macrophage; E: erythroid; GEMM: granulocyte-erythroid-macrophage-megakaryocyte; BFU-E: burst-forming unit-erythroid.

### Scalable System Supports Large-scale Erythroid Differentiation and Terminal Maturation

In previous studies, we reported a feeder-free static suspension condition for the generation of matured erythrocytes from the single cells of EB day 13 through two continuous steps of erythroid differentiation (ED) and terminal maturation (TM)^10^. Here we report the enhancement of the scalability of these two steps by applying dynamic suspension culture in scalable conditions (SF or TR, see the illustration in Fig. 2a). Considering that >70% of the suspension cells were CD34^+^ cells, we found that it was not necessary to isolate CD34^+^ cells for downstream erythroid production under a culture condition selectively favoring erythroid cell proliferation and differentiation. After EB formation and initial hematopoietic differentiation (Fig. 2a), single cells were harvested at day 13 by passing the entire culture through a 40-μm cell strainer and immediately transfer to the following differentiation steps. This simple “filter-and-go” process would be highly favorable for the scale-up bioprocess. Previous studies found that even mild agitation at 15-20 rpm in stirred tank bioreactor had significant impact on the *ex vivo* erythropoiesis of peripheral blood CD34^+^ derived cultures^43^. We also confirmed that shear stress generated by agitation at medium speed range (40-50 rpm) in spinner flasks impeded the expansion rate of erythroid expansion and differentiation (Fig. 4a). By reducing the agitation to low speed range in SF (22.5 rpm) or TR at low rocking frequency (10-15 rpm), expansion in erythroid differentiation and terminal maturation were recovered and comparable as in the static condition, calculated at 211±22 fold and 237±56 fold increase, respectively (Fig. 4a).

**Figure 4.**
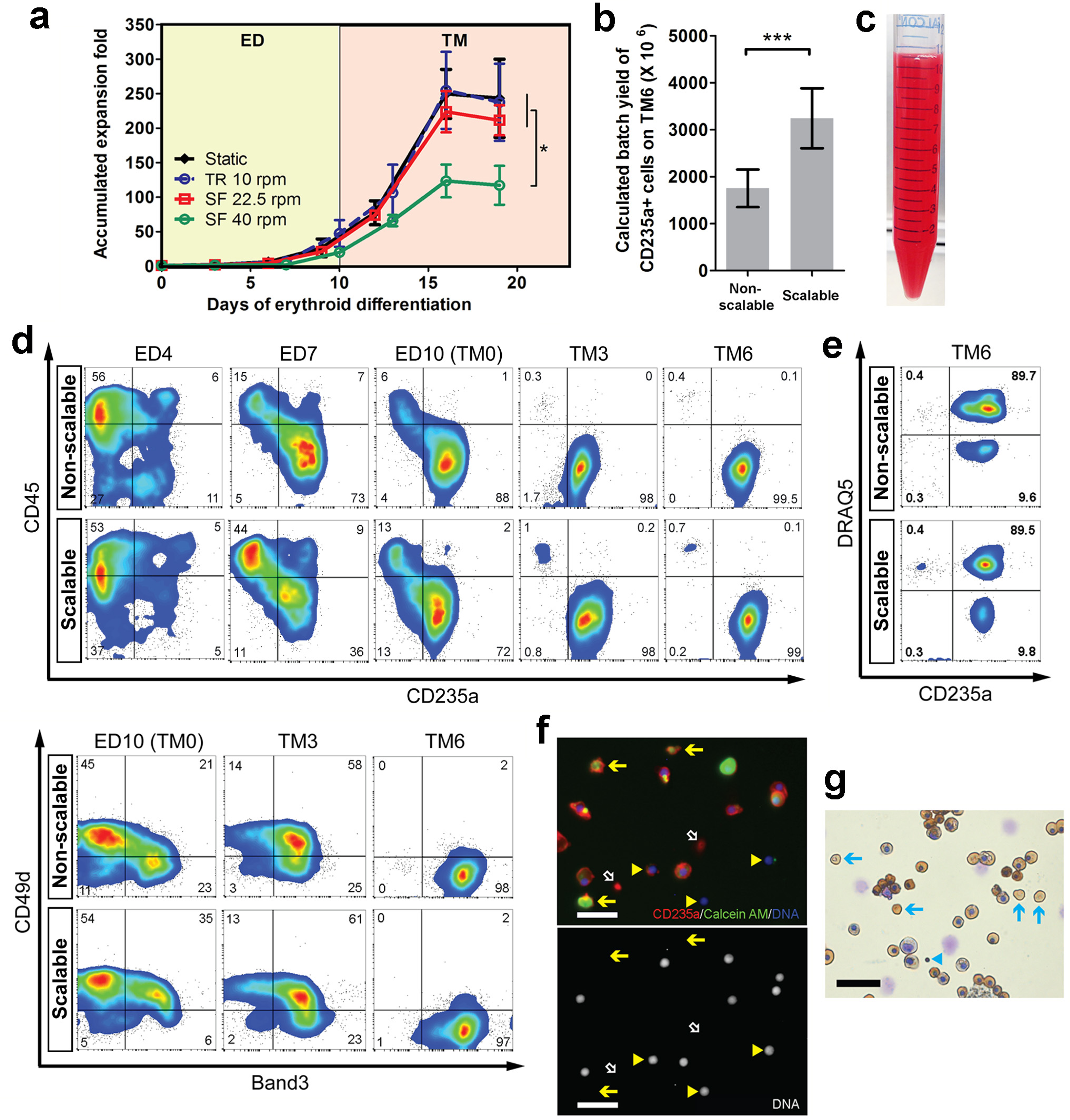
A scalable process of erythroid differentiation and terminal maturation. (a) Optimization of the agitating speed in scalable vessels for erythroid differentiation and terminal maturation (ED-TM) using the suspension cells from SF-TS-TR by analyzing accumulated expansion fold increase (n = 3 independent experiments, mean ± s.d.). All of the other conditions were significantly higher than SF 40 rpm condition (*P* < 0.05). (b) Calculated batch yield of CD235^+^ erythrocytes on day 6 of terminal maturation (TM6) from non-scalable process (forced aggregation EB and ED-TM in static suspension condition) and scalable process (SF-TS-TR EB and ED-TM in dynamic suspension condition on rocker at 10 rpm and/or in spinner flasks at 22.5 rpm (n = 4 independent experiments, mean ± s.d.). (c) Photo image of 1 × 10^9^ TM6 cells from the scalable process in 10 ml PBS. (d) Representative flow cytometry data of CD45, CD235a, CD49d and Band3 reveals the progressive changes of phenotype during ED-TM in both non-scalable and scalable process. All gates were set based on corresponding isotype controls. (e) Representative flow cytometry analysis of enucleation rate of TM6 cells using DRAQ5 staining for nuclei. All gates were set based on corresponding isotype controls. (f) Representative fluorescent images of TM6 cells generated from the scalable process. Solid arrows indicate live and metabolic active enucleated erythrocytes that were double stained by CD235a and Calcein AM but lack of nuclei and DRAQ5-staining of DNA. Triangle arrowheads indicate separated nuclei after enucleation, known as pyrenocytes. White open arrows indicate pieces of broken cell membrane that were only stained by CD235a (scale bar = 20 μm). (g) Wright-Giemsa (for cytoplasm and nuclear DNA) and benzidine (for hemoglobin) staining of TM6 cells from the scalable process. Most of the nucleated cells showed morphology of orthochromatic erythroblasts that contained hemoglobin (brown color by benzidine staining). Arrows indicate enucleated erythrocytes, and Triangle arrowheads indicate ejected nuclei (scale bar = 20 μm).

We compared the yield of erythrocytes from the process in scalable system (using SF-TS-TR for EB differentiation and SF or TR for ED-TM steps) and non-scalable system (using FA in 96-well plates for EB differentiation and static suspension culture for ED-TM steps^10^). The scalable process yielded 3.2±0.6×10^9^ CD235a^+^ erythrocytes, 1.5-fold higher than those from the non-scalable process (Fig. 4b). After scale-up to 1-litre spinner flasks, we achieved an ultra-high yield of 4.1×10^9^ hiPSC-derived erythrocytes starting from a 40-ml SF-TS-TR batch of EB differentiation. Taking these steps together, we established a complete procedure for the scalable production of human erythrocytes (Fig. 2a). A cell suspension containing 1×10^9^ cells in 10 ml PBS showed red color like diluted blood (Fig. 4c).

Phenotypic changes in the suspension cells from SF-TS-TR during the scalable ED-TM steps were tracked by flow cytometry (Fig. 4d and Supplementary Fig. S4). Plots of forward scatter and side scatter showed that the erythroblasts shrunk in size during the maturation period (Supplementary Fig. S4a). The gradual loss of CD45 and gain of CD235a expression indicated the loss of multipotent HSPCs and myeloid cell populations, and enrichment of erythroid population (Fig. 4d). When we analyzed the expression patterns of two erythroid stage-specific markers Band3 and CD49d (alpha4 integrin), we also observed a rapid and synchronized erythroblast maturation process from CD49d^+^Band3^-^, to CD49d^+^Band3^+^, and then to CD49d^-^Band3^+^, in line of previous studies using primary CD34^+^ cells or hiPSCs under the FA condition^10, 44^. After 6 days in TM culture, more than 95% of the cells were CD45^-^CD235a^+^Band3^+^CD71^+^. Only a few cells (0.5%-1.5%) expressed markers of myeloid lineages such as CD14, CD41, CD42a, CD15 and CD33 (Supplementary Fig. S4b) and no cells express lymphoid markers (data not shown). These data collected a highly pure population of erythrocytes as early as day 6 of TM. Enucleation (removal of nuclei) rate was measured by staining DNA with DRAQ5, followed by flow cytometry (Fig. 4e). Erythrocytes from SF-TS-TR showed 2.8%-11.1% of enucleation rates from multiple experiments, similar to the same batch of FA control group and the range of variation. The enucleated cells remained metabolically active and with intact cell membrane (Fig. 4f) and expressed hemoglobin protein (Fig. 4g).

### Scalable Strategy Supports Robust and Reproducible Generation of Erythrocytes from Multiple hiPSC Lines

In order to test the robustness and reproducibility of the scalable differentiation system, we repeated the process in two other hiPSC lines (TNC1 and E2) using the same SF-TS-TR protocol. TNC1 was derived from human erythroblasts that were established from the peripheral blood mononuclear cells of a sickle cell disease patient, while E2 was derived from human mesenchymal stem cells that were established from the bone marrow mononuclear cells of a healthy donor. Both hiPSC lines accomplished efficient and high-yield generation of HSPCs, erythroblasts and erythrocytes (Table 1 and Supplementary Fig. S5), confirming the robustness and adaptability of the scalable differentiation system.

**Table 1.** Summary of the yield in three hiPSC lines at the end of each differentiation steps from the scalable process

| hiPSC Lines | Yield ($\times 10^6$) from 40-ml batches of SF-TS-TR EB (n = 3, mean ± s.d.) | | | |
| --- | --- | --- | --- | --- |
|  | EB13<br>CD34<sup>+</sup>CD45<sup>+</sup> | ED10<br>CD36<sup>+</sup>CD235a<sup>+</sup> | TM6<br>CD235a<sup>+</sup> | Enucleated |
| **BC1** | 17 ± 5 | 843 ± 366 | 3947 ± 418 | 385 ± 206 |
| **E2** | 12 ± 4 | 653 ± 82 | 3928 ± 1017 | 206 ± 161 |
| **TNC1** | 17 ± 4 | 1096 ± 331 | 5942 ± 2002 | 564 ± 192 |

### Erythrocytes Generated from the Scalable System Show Adult Hemoglobin Expression and Oxygen Carrying Function

We analyzed the expression of hemoglobin genes particularly β-globin gene at both mRNA and protein levels. The mRNA level increased dramatically along the differentiation. On TM day 7, cells from SF-TS-TR showed similar mRNA level of *HBB* and *HBG* as erythrocytes derived from cord blood CD34^+^ cells that were induced ED-TM in the same medium and scalable settings (Fig. 5a). We further found that ~64% of TM day 7 cells from SF-TS-TR contained β-globin, while nearly 100% of the cells expressed γ-globin (Fig. 5b). These cells produced β-globin in the similar level as those from FA (Fig. 5b-c).

**Figure 5.**
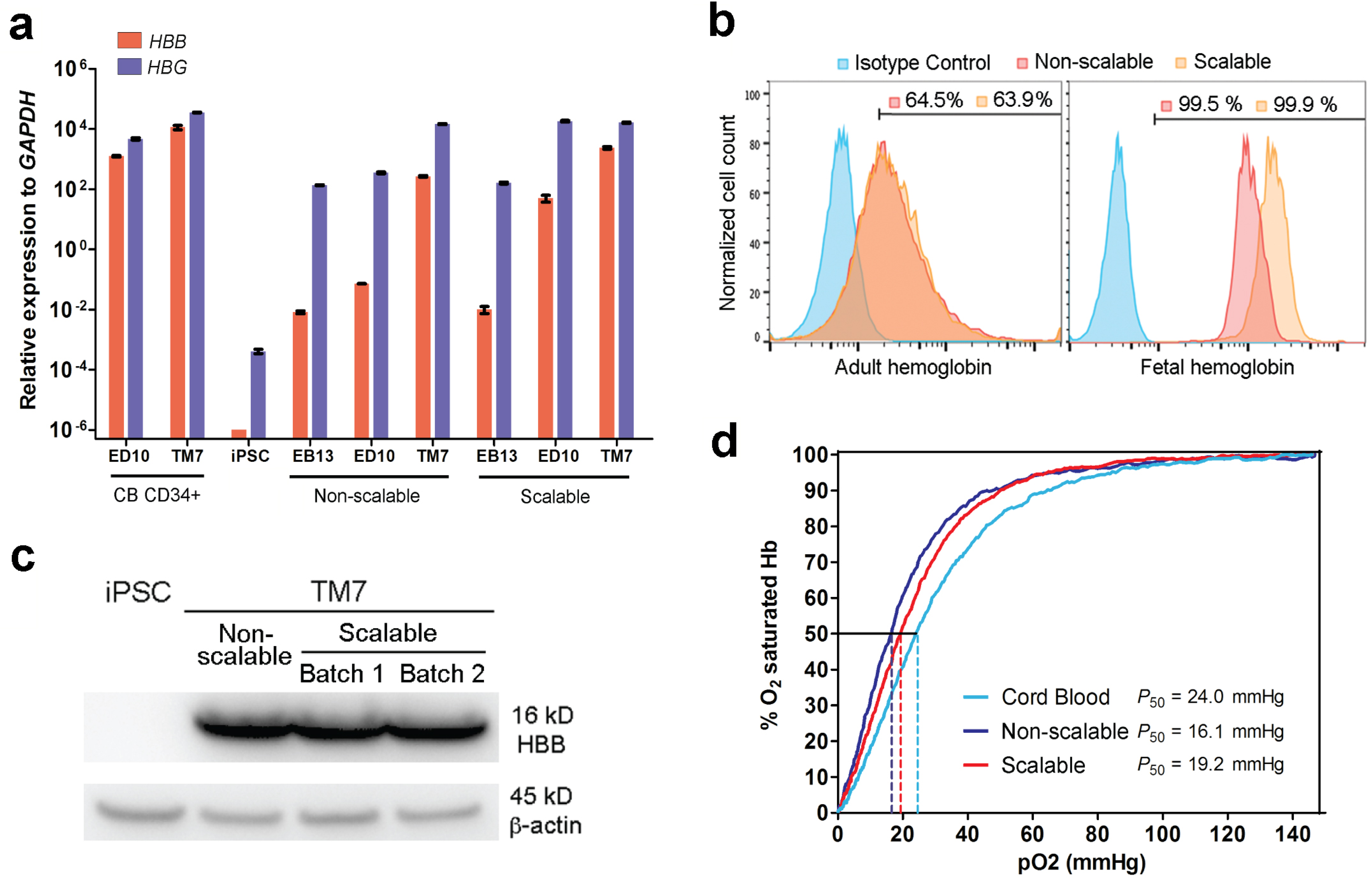
Hemoglobin expression and oxygen binding function of iPSC-derived erythrocytes from scalable production. (a) Quantitative RT-PCR analysis of *HBB* and *HBG* gene expression in the cells generated from non-scalable process (forced aggregation EB and ED-TM in static suspension condition) and scalable process (SF-TS-TR EB and ED-TM in dynamic suspension condition) at different stages, including undifferentiated iPSCs, total suspension cells after EB differentiation day 13 (EB13), erythroid differentiation day 10 (ED10) and terminal maturation day 7 (TM7). Erythrocytes derived from cord blood CD34^+^ cells (CB CD34^+^) following the scalable ED-TM in dynamic suspension conditions were analyzed together as a control (normalized to endogenous GAPDH as a housekeeping control; n = 2 independent experiments with 3 technical replicates each, mean ± s.d.). (b) Representative flow cytometry plots of intracellular adult and fetal hemoglobin protein in TM7 cells. (c) Western blot analysis of human adult β-globin (HBB) protein. BC1 iPSCs and TM7 cells from non-scalable process and two independent runs of scalable process were loaded as 10 μg total protein. β-actin was stained as a housekeeping control. (d) Oxygen-hemoglobin dissociation curves of TM7 cells derived from non-scalable and scalable process measured by a Hemox Analyzer, using primary cord blood red blood cells as a control.

We also determined whether the hemoglobin in the cells was functional to bind to O_2_ molecules at high concentration and release them at low concentration. Oxygen-hemoglobin dissociation curve of BC1-derived erythrocytes in scalable settings were measured and found to be similar to the curve of primary cord blood RBCs (Fig. 5d). Interestingly, TM day 6 cells from SF-TS-TR method (*P*_50_ = 19.2 mmHg) showed lower O_2_ binding affinity than those from FA method (*P*_50_ = 16.1 mmHg), and were closer to cord blood (*P*_50_ = 24.0 mmHg). It is known that fetal hemoglobin has a higher affinity to bind to O_2_ and that cord blood express both fetal and adult hemoglobin. This result is consistent with the data that the content of fetal hemoglobin in the erythrocytes from SF-TS-TR was lower than erythrocytes from FA method, but still higher than in cord blood.

## Discussion

Along with the flourish of transformative hiPSC technologies, cell culture conditions for scalable expansion of undifferentiated hiPSCs have been improved significantly^37, 45-47^. Chemically defined, feeder-and xeno-free culture media and substrates are able to support long-term and scalable expansion of hiPSCs while keeping a normal karyotype as we have demonstrated^48^. These improvements suggest a bypass to the elusive goal of *ex vivo* expansion of human HSPCs, especially long-term stem cell activities, which remains challenging^3^. In this way, one can expand undifferentiated hiPSCs first into a large number followed by a large-scale and high-efficiency differentiation process of HSPC production, to achieve a high yield of a final blood cell product such as erythrocytes. During this multi-step differentiation process, the generation of HSPCs from hiPSCs that are able to generate multiple types of mature hematopoietic cells is the rate-limiting step^14^. In the current study, we demonstrated the first complete procedure for scalable production of erythrocytes from hiPSCs under a feeder-free and xeno-free culture condition, with an innovative EB differentiation step.

The current success was based on our previous study where we demonstrated that hiPSCs can form cell spheres of an average equivalent diameter of ~200 μm in spinner flask culture under an optimized speed in E8 medium^36^. Each sphere contains approximately 3,000 cells, which correspond to the optimized number of input hiPSCs to form one EB per well in the FA EB formation method. For the EB formation in suspension in differentiation medium, we found that cell viability on day 1 was a critical checkpoint for differentiation efficiency, because apoptotic or dead cells wrapped inside the EBs would lead to break or failure of EB formation. hiPSCs seeded as single cells on day 0 were most vulnerable to shear-induced cell death, whereas insufficient shear would cause agglomeration and lead to transfer deficiency of O_2_, nutrients and metabolic wastes. Therefore, we sought to find a balance by choosing the appropriate platform and optimizing the agitating speed and culture medium formulation for EB formation in the critical first two days. We also found that different hiPSC lines had distinct sensitivity and tolerance to shear stress in spinner flasks, indicated by cell survival and EB sizes. Some iPSC lines such as BC1 and E2 survived poorly in spinner culture with the SFM-EB medium, even in slow agitation. To overcome this, we reported here that a new EB medium (based on the original E8 medium used for expanding hiPSCs) significantly enhanced the cell survival in spinner flasks. Moreover, we observed that physiological O_2_ tension (5%) that was known to enhance hiPSC survival also enhanced EB formation and differentiation, using an oxygen monitoring and controlling system (unpublished data, 2014-2015). It was reported that the dissolved O_2_ level in the media affected the rate and maturation of erythroblasts *in vitro*^49^. This oxygen monitoring and controlling system would also provide a suitable apparatus for further optimization of erythrocyte terminal differentiation by tuning O_2_ tension in the culture.

Shear stress induced by agitation is also known to play an important role in iPSC differentiation^50, 51^. The glass-ball type spinner flask produces laminar flow (*Re* < 1000) and mild shear rate less than 0.15 N/m^2^ ^36^, while tissue culture flasks on a rocker mimics a wave-type bag bioreactor that generates much lower shear of approximately 0.01 N/m^2^ measured in previous studies^52^. A static condition causes no shear on EBs. Compared to the freely floating EBs in a continuous suspension condition, the stepwise SF-TS-TR procedure showed higher differentiation efficiency and reproducibility. The enhancement may be due to the static condition and the loose attachment of EBs to the surface of cell culture vessels. This semi-adhesion might provide better environment for the generation of hemogenic endothelial cells and strengthened the up-regulation of many critical genes related to mesodermal and hematopoietic development such as *KDR*, *GATA2*, *RUNX1*, *HHEX* and *MYB*^39^. The mild agitation on a rocker in the following step might be beneficial for the hematopoietic differentiation by enhancing transport and preventing complete attachment leading to loss of EB structures. The CD34^+^CD45^+^ cells generated from the SF-TS-TR procedure appear to be definitive hematopoietic progenitor cells because they can form multiple types of myeloid cells and erythrocytes that express HBB and mature to enucleated cells. The suspension cells harvested at EB day 13 enriched for CD34^+^CD45^+^ cells contained few CD235a^+^ cells before EB day 13, the early onset of which was suggested to be an indicator of primitive hematopoiesis independent of CD34^+^CD45^+^ HSPC formation^53^.

The SF-TS-TR method requires 8.3-fold more input of hiPSCs per run than the FA method to initiate EB differentiation in current settings, which causes about 3-fold decrease in the efficiency of CD235a^+^ erythrocyte generation per input hiPSC on TM6. However, the batch yield is still about 2-fold higher. More importantly, the scalability of the SF-TS-TR system enables a volumetric scale-up process using 100-litre to 500-litre stirred tank bioreactors and rocking bag bioreactors, which is more preferable in the manufacture concept other than the linear scale-out process of using the FA method. By one batch of hiPSC differentiation, we generated 4.1×10^9^ CD235a^+^ erythrocytes using 1-litre spinner flasks at a density of approximately 4×10^6^ cells/ml. Therefore, it is possible to produce sufficient erythrocytes for a clinical unit of blood (containing 2×10^12^ RBCs) from one batch of 500-litre stirred tank bioreactor or WAVE bag bioreactor. Although reducing the cost remains a challenge of *ex vivo* generation of transfusable RBCs from hiPSCs^54^, the concept and improvement demonstrated in our study is an important step forward leading to a feasible solution of reducing manufacturing cost by expanding production scale. Substantial improvement of medium that can support higher differentiation efficiency and culture density would be very helpful to further manufacturing purpose.

The HSPCs generated from this scalable system were confirmed to exhibit comparable phenotype and potential for multi-lineage differentiation as the HSPCs from the FA method that were extensively evaluated in the previous studies^10^. Despite the substantially increased contents of functional hemoglobin, the enucleation rate (2%-15%) we achieved so far was still low as compared to those derived from primary CD34^+^CD45^+^ cells from cord blood and bone marrow. The enucleation rate also varied among hiPSC lines (and from batch-to-batch to a less degree), which is a shared problem among other studies on hiPSC-derived erythrocytes^9, 12, 55^. Although some studies suggested that the erythroblasts generated *in vitro* could eventually enucleate in circulation when being injected *in vivo*^55^, it would be ideal to reach full enucleation and eliminate all nucleated cells before being transfused into recipients. More efforts are needed to further improve the enucleation of hiPSC-derived erythrocytes. Recent reports on transcription profile of enucleating erythroblasts derived from cord blood CD34^+^ cells^44^ and hESCs^56^ provide an insight in the molecular signature of the enucleation process and may lead us to find more efficacious solutions to improve enucleation in suspension culture conditions.

In summary, we generated erythrocytes from three hiPSC lines in a scalable differentiation system that combined spinner flask and rocker for EB differentiation. We achieved high batch yield, β-globin expression and complete terminal maturation in the hiPSC-derived erythrocytes. The ease of operation, the simplicity of cytokine combination and the high level of chemical definition of the components added favorable features to this system in a scale-up point of view. Furthermore, because all the components used in our system for the entire expansion and differentiation process are formulated with recombinant human proteins, Food and Drug Administration (FDA)-approved biologics, or materials from human sources, this xeno-free and cGMP-compliant system provides the opportunity to produce clinical-grade erythrocytes from hiPSCs in a large-scale bioprocess for therapeutic purposes. Therefore, this study is a significant step forward towards the goal of large-scale generation of RBCs for therapeutic purposes.

## Methods

### EB Differentiation Using SF-TS-TR Method

BC1 and TNC1 hiPSCs were expanded in 100-ml spinner flasks (CELLSPIN, Integra Biosciences) as cell aggregates in E8 medium (Essential 8, Thermo Fisher Scientific) as previously described^36^ to yield a large quantity. Cell aggregates were dissociated into single cells by Accutase and inoculated at 4 × 10^5^ cells/ml in spinner flasks at agitating speed of 35-50 rpm to initiate EB formation in 40 ml E8-based EB-formation medium (E8-EB medium), which is E8 medium supplemented with 10 ng/ml BMP4 (R&D Systems), 0.5% w/v human albumin (Plasbumin-25, Amgen), and 10 μM ROCK inhibitor Y27632 (Stemgent). This was marked as day 0 of differentiation. On day 1, stop agitation, settle the EBs at the bottom of the flask, and change ~70% of the medium with fresh E8-EB medium to remove dead cells. All medium was replaced by SFM-based EB differentiation medium^30^ on day 2. On day 3, we transferred the culture into a 150-cm^2^ tissue culture flask, and placed in static (TS). After changing medium on day 6, we put the tissue culture flask on a platform rocker at a frequency of 10-15 rpm (TR). Then we changed ~70% of the medium on day 8, day 10, and day 13. Nplate (Amgen) were added at a final concentration of 20 ng/ml in EB differentiation medium from day 11 to day 13 to enhance HSPC expansion. Single suspension cells were harvested on day 13 and sequentially cultured in erythroid differentiation medium that was also made xeno-free by replacing bovine serum albumin (BSA) with human albumin, with the same concentration of cytokines and holo-transferrin for 10 days as previously described^10^, followed by terminal maturation for 6-8 days^10^, in either TR or SF condition with low agitating speed. FA method (so called “spin-EB”) was applied as positive control as previously described^30^. When calculating yield, one batch of FA control was set as 8 plates of 96-well plates for the same 40-ml initial medium volume.

### Erythroid Differentiation and Terminal Maturation

Suspension cells harvested from EB day 13 were directly transferred into an erythroid liquid culture in tissue culture plates in static condition, on rocker, or in spinner flasks without the need of CD34^+^ isolation. The medium formulations for erythroid differentiation and terminal maturation were described previously^10, 31^, except that the BSA in SFM was replaced by human albumin to make it xeno-free^30^.

Additional methods can be found in the Supplementary Information.

### Statistics

All analysis of cell number, cell viability, EB size, and flow cytometry were collected from at least three independent experiments (n ≥ 3). Real-time quantitative PCR analysis was performed in two independent experiments with triplicate technical replicates each. The unpaired two-tailed Student’s t-test or two-way analysis of variance (two-way ANOVA) followed by Bonferroni’s posttest was done using GraphPad Prism 5 for the comparison between two groups or multiple groups, respectively. Data are presented as mean ± s.d. if not specified. Significance level was assigned as not significant (n.s.) *P* > 0.05, * *P* < 0.05, ** *P* < 0.01, and *** *P* < 0.001.

## Supplementary Information

Supplemental Information includes Supplemental Experimental Procedures and four Supplemental Figures and can be found with this article online.

## Acknowledgement

The authors thank Dr. Xiuli An at NYBC for providing the Band-3 antibodies, and Drs. Hao Bai and Yanfeng Liu at JHMI for providing PCR primers and scientific discussions. We also thank Laurel Mendelsohn at NHLBI for conducting O_2_ binding experiments using a Hemox-Analyzer. This work was supported in part by grants from Maryland State Stem Cell Research Fund (2011-MSCRFII-0088 and 2011-MSCRFE-0087) and from NIH (2R01 HL073781, R01 HL130676, U01 HL107446 and T32 HL007525). L.C. is also supported by Edythe Harris Lucas and Clara Lucas Lynn Chair in Hematology of Johns Hopkins University.

## Author Contribution Statement

Y.W., designed and performed research, collected and analyzed data, and wrote the manuscript. Y.G. and C.H. performed research. Z.Y., analyzed data. S.G., designed research, analyzed and interpreted data, and wrote the manuscript. L.C., designed research, analyzed and interpreted data, wrote the manuscript, and provided financial support.

## Additional Information

The authors indicate no potential conflicts of interest.

## References

1. Anstee, D.J. The relationship between blood groups and disease. Blood 115, 4635–4643 (2010).

2. Giarratana, M.C. et al. Proof of principle for transfusion of in vitro-generated red blood cells. Blood 118, 5071–5079 (2011).

3. Dahlberg, A., Delaney, C. & Bernstein, I.D. Ex vivo expansion of human hematopoietic stem and progenitor cells. Blood 117, 6083–6090 (2011).

4. Robinton, D.A. & Daley, G.Q. The promise of induced pluripotent stem cells in research and therapy. Nature 481, 295–305 (2012).

5. Kaufman, D.S. Toward clinical therapies using hematopoietic cells derived from human pluripotent stem cells. Blood 114, 3513–3523 (2009).

6. Peyrard, T. et al. Banking of pluripotent adult stem cells as an unlimited source for red blood cell production: potential applications for alloimmunized patients and rare blood challenges. Transfusion medicine reviews 25, 206–216 (2011).

7. Migliaccio, A.R., Whitsett, C., Papayannopoulou, T. & Sadelain, M. The potential of stem cells as an in vitro source of red blood cells for transfusion. Cell stem cell 10, 115–119 (2012).

8. Shah, S., Huang, X. & Cheng, L. Concise review: stem cell-based approaches to red blood cell production for transfusion. Stem cells translational medicine 3, 346–355 (2014).

9. Dias, J. et al. Generation of red blood cells from human induced pluripotent stem cells. Stem cells and development 20, 1639–1647 (2011).

10. Huang, X. et al. Production of Gene-Corrected Adult Beta Globin Protein in Human Erythrocytes Differentiated from Patient iPSCs After Genome Editing of the Sickle Point Mutation. Stem cells 33, 1470–1479 (2015).

11. Kobari, L. et al. Human induced pluripotent stem cells can reach complete terminal maturation: in vivo and in vitro evidence in the erythropoietic differentiation model. Haematologica 97, 1795–1803 (2012).

12. Lapillonne, H. et al. Red blood cell generation from human induced pluripotent stem cells: perspectives for transfusion medicine. Haematologica 95, 1651–1659 (2010).

13. Yang, C.T. et al. Human induced pluripotent stem cell derived erythroblasts can undergo definitive erythropoiesis and co-express gamma and beta globins. British journal of haematology 166, 435–448 (2014).

14. Slukvin, II Hematopoietic specification from human pluripotent stem cells: current advances and challenges toward de novo generation of hematopoietic stem cells. Blood 122, 4035–4046 (2013).

15. Lu, S.J. et al. Biologic properties and enucleation of red blood cells from human embryonic stem cells. Blood 112, 4475–4484 (2008).

16. Choi, K.D., Vodyanik, M. & Slukvin, II Hematopoietic differentiation and production of mature myeloid cells from human pluripotent stem cells. Nature protocols 6, 296–313 (2011).

17. Ledran, M.H. et al. Efficient hematopoietic differentiation of human embryonic stem cells on stromal cells derived from hematopoietic niches. Cell stem cell 3, 85–98 (2008).

18. Rafii, S. et al. Human ESC-derived hemogenic endothelial cells undergo distinct waves of endothelial to hematopoietic transition. Blood 121, 770–780 (2013).

19. Qiu, C., Olivier, E.N., Velho, M. & Bouhassira, E.E. Globin switches in yolk sac-like primitive and fetal-like definitive red blood cells produced from human embryonic stem cells. Blood 111, 2400–2408 (2008).

20. Uenishi, G. et al. Tenascin C promotes hematoendothelial development and T lymphoid commitment from human pluripotent stem cells in chemically defined conditions. Stem cell reports 3, 1073–1084 (2014).

21. Itskovitz-Eldor, J. et al. Differentiation of human embryonic stem cells into embryoid bodies compromising the three embryonic germ layers. Molecular medicine 6, 88–95 (2000).

22. Kennedy, M., D’Souza, S.L., Lynch-Kattman, M., Schwantz, S. & Keller, G. Development of the hemangioblast defines the onset of hematopoiesis in human ES cell differentiation cultures. Blood 109, 2679–2687 (2007).

23. Zambidis, E.T., Peault, B., Park, T.S., Bunz, F. & Civin, C.I. Hematopoietic differentiation of human embryonic stem cells progresses through sequential hematoendothelial, primitive, and definitive stages resembling human yolk sac development. Blood 106, 860–870 (2005).

24. Cerdan, C., Rouleau, A. & Bhatia, M. VEGF-A165 augments erythropoietic development from human embryonic stem cells. Blood 103, 2504–2512 (2004).

25. Chang, K.H. et al. Definitive-like erythroid cells derived from human embryonic stem cells coexpress high levels of embryonic and fetal globins with little or no adult globin. Blood 108, 1515–1523 (2006).

26. Lachmann, N. et al. Large-scale hematopoietic differentiation of human induced pluripotent stem cells provides granulocytes or macrophages for cell replacement therapies. Stem cell reports 4, 282–296 (2015).

27. Ye, Z. et al. Human-induced pluripotent stem cells from blood cells of healthy donors and patients with acquired blood disorders. Blood 114, 5473–5480 (2009).

28. Ng, E.S., Davis, R.P., Azzola, L., Stanley, E.G. & Elefanty, A.G. Forced aggregation of defined numbers of human embryonic stem cells into embryoid bodies fosters robust, reproducible hematopoietic differentiation. Blood 106, 1601–1603 (2005).

29. Ungrin, M.D., Joshi, C., Nica, A., Bauwens, C. & Zandstra, P.W. Reproducible, ultra high-throughput formation of multicellular organization from single cell suspension-derived human embryonic stem cell aggregates. PloS one 3, e1565 (2008).

30. Liu, Y. et al. Efficient generation of megakaryocytes from human induced pluripotent stem cells using food and drug administration-approved pharmacological reagents. Stem cells translational medicine 4, 309–319 (2015).

31. Ye, Z. et al. Differential sensitivity to JAK inhibitory drugs by isogenic human erythroblasts and hematopoietic progenitors generated from patient-specific induced pluripotent stem cells. Stem cells 32, 269–278 (2014).

32. Kempf, H. et al. Controlling expansion and cardiomyogenic differentiation of human pluripotent stem cells in scalable suspension culture. Stem cell reports 3, 1132–1146 (2014).

33. Correia, C. et al. Combining hypoxia and bioreactor hydrodynamics boosts induced pluripotent stem cell differentiation towards cardiomyocytes. Stem cell reviews 10, 786–801 (2014).

34. Ting, S., Chen, A., Reuveny, S. & Oh, S. An intermittent rocking platform for integrated expansion and differentiation of human pluripotent stem cells to cardiomyocytes in suspended microcarrier cultures. Stem cell research 13, 202–213 (2014).

35. Vosough, M. et al. Generation of functional hepatocyte-like cells from human pluripotent stem cells in a scalable suspension culture. Stem cells and development 22, 2693–2705 (2013).

36. Wang, Y. et al. Scalable expansion of human induced pluripotent stem cells in the defined xeno-free E8 medium under adherent and suspension culture conditions. Stem cell research 11, 1103–1116 (2013).

37. Kehoe, D.E., Jing, D., Lock, L.T. & Tzanakakis, E.S. Scalable stirred-suspension bioreactor culture of human pluripotent stem cells. Tissue engineering. Part A 16, 405–421 (2010).

38. Chou, B.K. et al. Efficient human iPS cell derivation by a non-integrating plasmid from blood cells with unique epigenetic and gene expression signatures. Cell research 21, 518–529 (2011).

39. Elcheva, I. et al. Direct induction of haematoendothelial programs in human pluripotent stem cells by transcriptional regulators. Nature communications 5, 4372 (2014).

40. Ran, D. et al. RUNX1a enhances hematopoietic lineage commitment from human embryonic stem cells and inducible pluripotent stem cells. Blood 121, 2882–2890 (2013).

41. Paz, H., Lynch, M.R., Bogue, C.W. & Gasson, J.C. The homeobox gene Hhex regulates the earliest stages of definitive hematopoiesis. Blood 116, 1254–1262 (2010).

42. Dai, G. et al. Over-expression of c-Myb increases the frequency of hemogenic precursors in the endothelial cell population. Genes to cells: devoted to molecular & cellular mechanisms 11, 859–870 (2006).

43. Boehm, D., Murphy, W.G. & Al-Rubeai, M. The effect of mild agitation on in vitro erythroid development. Journal of immunological methods 360, 20–29 (2010).

44. An, X. et al. Global transcriptome analyses of human and murine terminal erythroid differentiation. Blood 123, 3466–3477 (2014).

45. Amit, M. et al. Dynamic suspension culture for scalable expansion of undifferentiated human pluripotent stem cells. Nature protocols 6, 572–579 (2011).

46. Olmer, R. et al. Suspension culture of human pluripotent stem cells in controlled, stirred bioreactors. Tissue engineering. Part C, Methods 18, 772–784 (2012).

47. Zweigerdt, R., Olmer, R., Singh, H., Haverich, A. & Martin, U. Scalable expansion of human pluripotent stem cells in suspension culture. Nature protocols 6, 689–700 (2011).

48. Wang, Y., Cheng, L. & Gerecht, S. Efficient and scalable expansion of human pluripotent stem cells under clinically compliant settings: a view in 2013. Annals of biomedical engineering 42, 1357–1372 (2014).

49. Rogers, H.M. et al. Hypoxia alters progression of the erythroid program. Experimental hematology 36, 17–27 (2008).

50. Earls, J.K., Jin, S. & Ye, K. Mechanobiology of human pluripotent stem cells. Tissue engineering. Part B, Reviews 19, 420–430 (2013).

51. Fridley, K.M., Kinney, M.A. & McDevitt, T.C. Hydrodynamic modulation of pluripotent stem cells. Stem cell research & therapy 3, 45 (2012).

52. Oncul, A.A., Kalmbach, A., Genzel, Y., Reichl, U. & Thevenin, D. Characterization of flow conditions in 2 L and 20 L wave bioreactors using computational fluid dynamics. Biotechnology progress 26, 101–110 (2010).

53. Kennedy, M. et al. T lymphocyte potential marks the emergence of definitive hematopoietic progenitors in human pluripotent stem cell differentiation cultures. Cell reports 2, 1722–1735 (2012).

54. Zeuner, A. et al. Concise review: stem cell-derived erythrocytes as upcoming players in blood transfusion. Stem cells 30, 1587–1596 (2012).

55. Hirose, S. et al. Immortalization of erythroblasts by c-MYC and BCL-XL enables large-scale erythrocyte production from human pluripotent stem cells. Stem cell reports 1, 499–508 (2013).

56. Rouzbeh, S. et al. Molecular Signature of Erythroblast Enucleation in Human Embryonic Stem Cells. Stem cells (2015).

